# Optimal foraging requires coordinating decisions with movements

**DOI:** 10.1101/2025.09.19.677254

**Authors:** Clara Saleri, Gislène Gardechaux, Alexandre Foncelle, David Thura

## Abstract

Decision-making and motor execution are both constrained by time and energy, and their optimization depends on cost-benefit trade-offs. While many studies suggest that decisions and movements follow common economic principles and can be jointly regulated to maximize reward rate, recent findings indicate that these two processes can also be decoupled or modulated in compensatory ways. This variability raises the question of whether decision-action coordination reflects an optimal strategy aimed at maximizing reward rate, or rather a flexible but suboptimal adaptation to contextual constraints. To address this issue, we trained three macaque monkeys to perform a reaching-based foraging task. The observed adjustments in decision-action coordination of the three monkeys are consistent with the pursuit of a strategy that approximates the theoretical optimum of reward intake rate at the single trial level. The present findings indicate, both behaviorally and theoretically, that coordination between decision and movement is required to optimize behavior in economic terms.

## Introduction

During decision-making, the probability of making the right choice and the time available for that choice must be considered in light of the context and objective in which the decision is made. When time is not a constraint, an individual may seek to optimize the accuracy of their decisions. Very often, however, decisions are subject to time pressure (Chittka et al., 2009). The impact of time on decision-making occurs in several ways. First, in urgent situations, individuals must make a decision before the fixed time runs out, even if they have not fully deliberated. Imagine for instance a driver reacting to a sudden obstacle on the road. She has to choose how to avoid it really fast (Reddi and Carpenter, 2000; Cisek et al., 2009). Moreover, long deliberations involve an opportunity cost, because while you are investing time doing something, you cannot be doing something else that might be more rewarding (Payne et al., 1996; Rieskamp and Hoffrage, 2008). Finally, even in the absence of explicit deadlines or other opportunities, time by itself diminishes the value of the reward or success made possible by informed decision-making (Myerson and Green, 1995; Green et al., 1999).

The natural way to consider choice accuracy while limiting the impact of time is to trade-off decision speed against accuracy (see Heitz, 2014 for a review). This strategy predicts that individuals will seek to maximize not decision accuracy, but the *rate* of correct decisions. In experimental paradigms in which participants are faced with dozens of successive decisions to make, studies indeed indicate that human subjects behave in ways that optimize their success rate rather than the accuracy of their decisions (Bogacz et al., 2010; Balci et al., 2011).

During interactive behavior, decisions alone do not lead to rewards. Decisions are always expressed through movements, and it is through these actions that rewards can eventually be obtained (Shadlen et al., 2008; Cisek and Kalaska, 2010; Gordon et al., 2021). Crucially, goal-directed movements appear to be governed by the same economic principles that determine decisions, including the trade-off between accuracy and time and between time and energy expenditure (or effort) (Shadmehr et al., 2019; Shadmehr and Ahmed, 2020).

Because the same economic principles, namely reward versus costs (whose balance is sometimes referred to utility), appear to govern both decisions and movements, it has been proposed that these two processes are jointly controlled, allowing for the optimization of the rate of success (Thura et al., 2014, 2025; Carland et al., 2019). This joint control, or co-regulation, of decisions and movements makes good ecological sense. For example, a teammate of a basketball player suddenly offering a passing opportunity will trigger a quick reaction from that player to execute a vigorous pass that will increase the probability of success given the rapid evolution of the opponents’ position.

However, control of decisions and actions appears to not be as simple as that. Recent studies in humans, monkeys, and rodents indeed indicate that decision and movement vigor can also be decoupled, and even controlled in opposite ways (Reynaud et al., 2020; Fievez et al., 2024; Morvan et al., 2024; Saleri and Thura, 2024; Leroy et al., 2025). Some evidence of decoupling comes from studies where subjects had to complete a given number of correct decisions in each block of a decision task with different motor constraints. When motor constraints required slow and accurate reaching (e.g., moving to a small target), decisions became faster whereas when movements could be fast, decisions slowed down. These results point to a compensatory relationship between decision-making and movement durations (Saleri Lunazzi et al., 2021, 2023; Leroy et al., 2025).

While the two identified coordination modes between decisions and movements, namely the “co-regulation” and the “compensation”, have been interpreted as context-dependent strategies underlying economic optimization principles, particularly the minimization of reward devaluation over time (Thura et al., 2025), this hypothesis suffers from a lack of more direct investigation.

The goal of the present study is to evaluate the importance of coordinating decisions with subsequent movements during natural behavior. More specifically, we aim to assess whether such coordination allows to approximate optimal behavior, in terms of reward rate, or whether it merely reflects flexible, yet suboptimal, adaptations.

To this aim, we trained three macaque monkeys to perform a reaching-based foraging task in which reward rhythm and movement cost vary between trials and blocks of trials, respectively. A foraging task is particularly well situated to address the goal mentioned above because it includes a succession of decisions about when to end an exploitation followed by movements executed to “explore” and start new ones; it is an ecological paradigm that reflects the reasons the primate brain evolved (Calhoun and Hayden, 2015); and it is a paradigm that is well described at the theoretical level, allowing to compare animals’ behavior with respect to optimal strategies derived from these theories (Charnov, 1976; Yoon et al., 2018).

In such a context where a subject has to make a succession of decisions to leave depleting sources of reward to explore other locations, when reward rhythm and movement cost vary between trials and blocks of trials, during multiple sessions, we make the hypothesis that coordinating decision duration with movement vigor (duration and/or speed) contributes to improve the intake rate. This hypothesis predicts that any improvement in the animals’ intake rate will be accompanied by an adjustment in the mode of coordination of decision and movement, and that this mode of coordination corresponds to the optimal strategy at the theoretical level with respect to the reward rate.

## Material and methods

### Subjects, ethical framework, and testing context

Experiments were conducted with three rhesus macaque monkeys (Macaca *mulatta* - monkey G, female, 9 kg, 11 years old, right-handed; monkey B, male, 6 kg, 5 years old, left-handed; monkey D, male, 6 kg, 5 years old, left-handed). Data were collected over a total of 84 sessions for monkey G, 50 for monkey B and 36 for monkey D.

Ethical permission was provided by “Comité d’Éthique Lyonnais pour les Neurosciences Expérimentales” (CELYNE), C2EA #42, ref: C2EA42-11-11–0402-004, and by the French Ministry of Research and Higher Education (initially obtained in 2020, renewed in 2024). Monkey housing and care were in accordance with the European Community Council Directive (2010) and the Weatherall report, “The use of non-human primates in research”. Laboratory authorization was provided by the “Préfet de la Région Rhône-Alpes” and the “Directeur départemental de la protection des populations” under Approval Number D69 029 06 01.

Behavioral data from monkey G were collected head-fixed and simultaneously with electrophysiological recordings which required the manipulation of microelectrodes to isolate neuronal activity (which will be the topic of future publications). To this end she was implanted with a titanium head fixation post and with one form-fitting recording chamber (Gray Matter Research). All surgical procedures were carried out under anesthesia and strict aseptic conditions. To ensure the animal’s well-being and reduce any stress associated with handling, light sedation was administered prior to each session using Domitor^Ⓡ^ (medetomidine) at a dose of 0.01mg/kg, by intramuscular injection. The behavioral task began once monkey G was fully awake and attentive, typically about 30-40 minutes after the administration of Domitor. By contrast, because monkeys B and D were not yet implanted for neurophysiological recordings, their behavioral data were acquired under head-free conditions and without the light sedation used for monkey G.

### Experimental apparatus

The monkeys sat in a primate chair and were trained to perform a simple foraging task based on planar arm movements. They held a lever in their dominant hand (right for monkey G, left for monkeys B and D) and a digitizing tablet (GTCO CalComp) continuously recorded its horizontal and vertical positions (100 Hz with 0.013 cm accuracy). The visual display was projected by a VIEWPixx monitor (VPixx Technologies; 120 Hz refresh rate) onto a mirror suspended 25 cm above and parallel to the digitizer plane (Figure 1A). Unconstrained eye movements and pupil area were recorded using an Eyelink 1000 (SR Research) infrared camera (data not shown).

**Figure 1.**
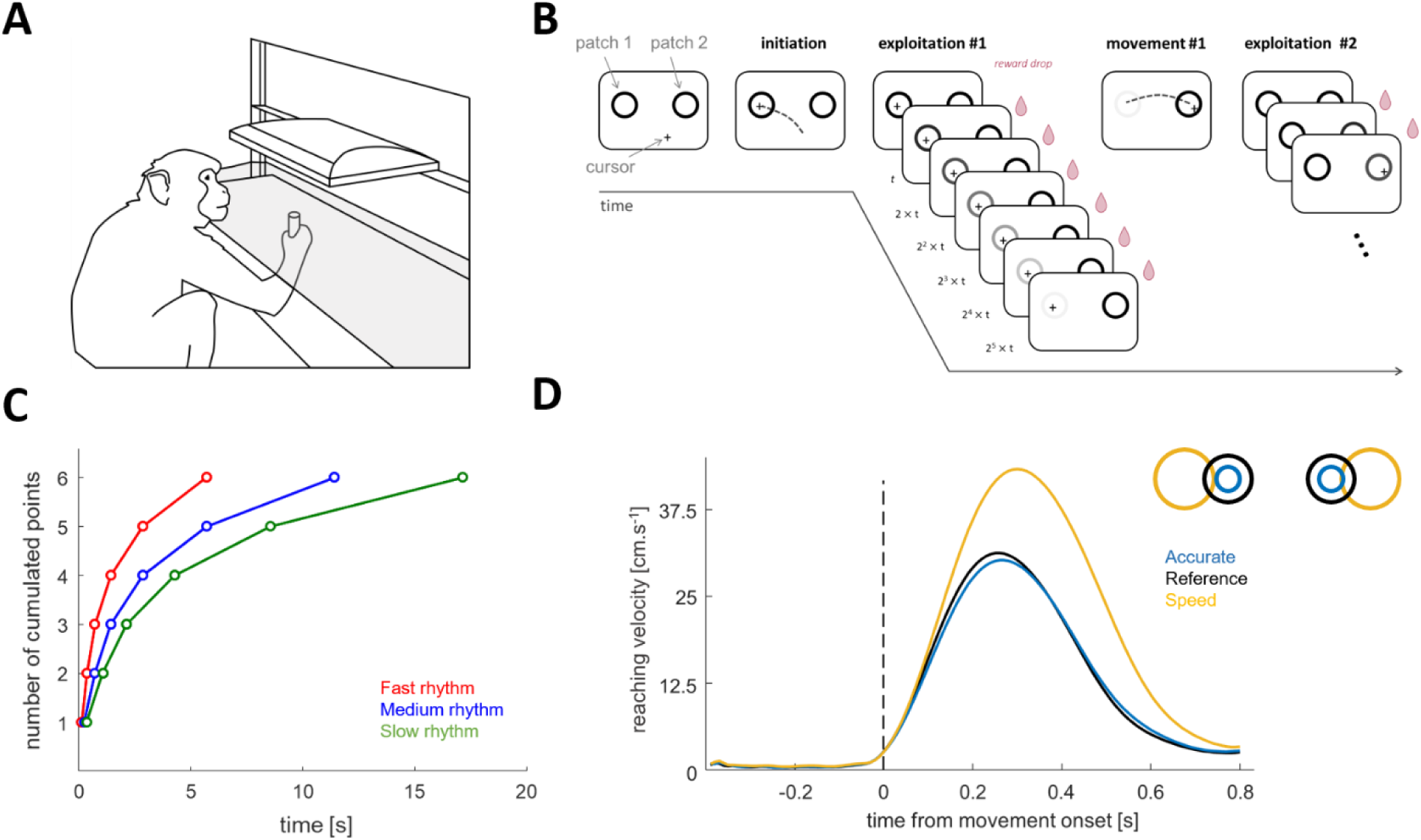
Experimental setup, design and conditions. **A**: Experimental apparatus. The monkeys sat in a primate chair and performed arm movements to control a lever held in their dominant hand. A digital tablet continuously recorded the horizontal and vertical positions of the lever. Visual stimuli and a cursor, which served as visual feedback of the lever, were projected onto a semi-transparent mirror positioned parallel above the digital tablet. **B**: Time course of a trial in the foraging task (see text for details). **C**: The three different reward rhythms, illustrating the timing of the six potential drops of juice that the monkey could obtain by keeping the cursor into a patch (slow in green, medium in blue and fast in red). **D**: Average velocity profiles of all reaching movements executed by monkey G in each of the three motor conditions (curves aligned to the onset of the movements; the same patterns were observed for monkey B and monkey D). Motor conditions are performed in distinct blocks of trials, representing the combinations of target size and distance (insert panel). The reference condition is illustrated in black, the accurate condition in blue and the speed condition in orange.

### Behavioral task

The three macaques have been trained to execute a computerized, reaching-based foraging task (Figure 1B) in the apparatus described above. In this task animals manipulated a lever and performed a reaching movement toward one of two possible visual targets (the reward sources, or “patches”) to collect drops of fruit juice (the reward). Patches were indicated by two circles placed horizontally. The position of the lever was materialized on the screen by a black cross (hereafter referred to as the ’cursor’). Once the lever was placed in a patch and held still for 100ms, juice drops were delivered at a rhythm that decreased over time, mimicking the exploitation of a depleting source of rewards. When the time interval between two drops was perceived to be too long, the monkeys could decide to leave the current patch by executing a reaching movement toward the other patch to start another exploitation. Reward rhythms and reaching movement costs were varied within and between blocks of trials, respectively (see the experimental conditions below).

From the experimenter’s point of view, each drop of fruit juice collected by the monkey corresponded to a point. At the start of each session the experimenter set a number of points that the monkeys had to collect from the two possible patches by alternating periods of static exploitations followed by arm movements executed to reach the other patch, in a back-and-forth manner (from the right patch to the left one, then from the left patch to the right one, and so on). On the top of the patches, a gauge that filled up as animals progressed in the task (in terms of accumulated points/rewards) was displayed. We assumed that efficient task performance involved animals adopting a strategy that maximized point accumulation while minimizing time spent in the experimental room.

At each exploitation phase, the monkeys could obtain up to 6 drops of juice, with the time interval between two being systematically multiplied by a factor of approximately two (Table 1). This implies that the longer they stayed in a patch to exploit it, the longer they had to wait to get the next drop of juice. To visually reinforce the impression of exhaustion of a current source of reward, the color of the exploited circle gradually faded after each collected reward, disappearing completely by the 6th reward (Figure 1B).

**Table 1.** Timings of the 6 potential points for each reward rhythm level and proportion of each rhythm level faced by each monkey across sessions.

| Reward rhythm level | Timings [s] |  |  |  |  |  | % for monkey G | % for monkey B | % for monkey D |
| --- | --- | --- | --- | --- | --- | --- | --- | --- | --- |
|  | #1 | #2 | #3 | #4 | #5 | #6 |  |  |  |
| Slow | 0.363 | 1.074 | 2.142 | 4.282 | 8.565 | 17.152 | 33% | 30% | 30% |
| Medium | 0.244 | 0.718 | 1.430 | 2.855 | 5.710 | 11.422 | 35% | 40% | 40% |
| Fast | 0.126 | 0.362 | 0.718 | 1.429 | 2.855 | 5.708 | 32% | 30% | 30% |

In most foraging scenarios, the decision to leave a reward source to explore other opportunities depends on the current exploitation as well as the cost of the next exploration (Charnov, 1976; Hayden et al., 2011; Calhoun and Hayden, 2015). Therefore, by convention, we defined as a trial in this task an exploitation phase in one patch followed by an arm movement to the other patch. This sequence of exploitation-movement was repeated by the monkeys as long as they were sufficiently motivated to perform it. If the gauge was full at the end of the session, they received, in addition to their liquid rewards accumulated during the session and their daily ration of accommodation, a “jackpot” reward, most often a whole fruit.

As expected in this task, the monkeys mostly performed successions of exploitation in one patch followed by an arm movement directed toward the other patch. However, sometimes and for various reasons, the constraints of the task were sufficiently weak to allow other behavioral patterns, such as exploitation of a patch, followed by a short movement to exit that patch and a return to it. Without formally prohibiting it, several measures were planned to discourage such behavior. Especially, if they did not cross the median distance separating the two patches and returned to the same patch, the value of the patch was not reset. If the animals crossed this median line but returned to the same patch in which they had just collected rewards, the value of this patch was reset but a short time penalty delaying the delivery of the first drop was applied. Only trials in which an exploitation in a given patch was followed by moving toward the other patch were included in the analyses described below.

Moreover, because monkeys were not always uniformly engaged in the task throughout the session, a trial was considered valid for specific analyses if at least one reward/point was collected in the exploited patch, if the exploitation phase lasted less than 19 seconds (since no more rewards could be obtained beyond this delay), and if the duration of the reaching movement ranged between 80ms and 10 seconds (to exclude outlier or aberrant movements). This represented 74%, 84% and 71% of the total trials for monkey G, monkey B and monkey D, respectively.

### Experimental conditions

In this foraging task, the exploitation of a patch is subject to a decreasing rhythm of rewards collected. As previously mentioned, the time interval between each of the 6 potential rewards increases, requiring the monkey to wait longer for the next one. The rhythm varied between 3 levels from trial to trial. For the *Slow* rhythm, 17.2 ± 0.02 seconds (mean ± SD) were needed to obtain the 6 drops of juice, 11.4 ± 0.04 s for the *Medium* rhythm and only 5.7 ± 0.02 s for the *Fast* rhythm (see Figure 1C and Table 1 for the timing parameters of each drop in each of the three rhythms). Importantly, the rhythm of the next patch was unknown by the monkeys until they started to exploit it.

The probability of exploiting a patch with a slow, medium, or fast rhythm was manipulated within specific blocks of trials referred to as “environments”. The impact of the environment on animals’ behavior is beyond the scope of the present study. Relevant to the study however is the proportion of each reward rhythm faced by each monkey across sessions, which is reported in Table 1.

To assess the impact of movement costs on animals’ exploitation and movement behavior, the size and the distance of the patches were varied in three blocks of trials. In the *Accurate* condition, the patches were the smallest (1.25cm radius) and placed at a distance of 7cm from each other, encouraging slower and more precise movements compared to a *Reference* condition in which the patch size was fixed at 2.25cm radius (but still located 7cm apart, Figure 1D - insert panel). By contrast, in the *Speed* condition, the size and distance (2.75cm radius and 10cm apart) encouraged the monkey to make more vigorous movements compared to the reference condition (Figure 1D). During each session, monkeys performed blocks of approximately 50 trials per motor condition, continuing until achieving the daily objective of points. For all sessions, the reference condition was always presented first, while the order of the speed and accurate conditions was counterbalanced across sessions.

In summary, monkeys performed the task under three levels of reward rhythm and three levels of motor cost, resulting in 9 possible combinations of experimental conditions.

### Data analysis

Because the monkeys manipulated a lever to exploit and move in the task, the horizontal and vertical lever position data were used to calculate the different variables of interest described below. The kinematics data were first filtered using a 15th-degree polynomial filter and then differentiated to obtain a velocity profile. The onset and offset of each movement were determined by applying a velocity threshold of 2.5cm/s.

The exploitation phase (i.e. when the monkeys hold the lever inside one of the two patches and receive drops of juice at a decreasing rate) ends with the animal’s decision to leave the current patch and move to the other one. Thus, exploitation duration (ED) was calculated as the time between the stabilization of the cursor inside the current patch and the onset of the following movement executed to reach the other patch. For the “exploration” phase of each trial (i.e., the reaching movement), the velocity peak (VP) was determined as the maximum value between onset and offset of the movement executed to leave the exploited patch. The effectiveness of points/rewards collection was quantified at the session level using the intake rate (IR), defined as the number of points/rewards collected per minute of foraging. This metric was calculated by dividing the total number of points accumulated over the session by the total duration of the session.

The monkeys completed multiple experimental sessions on different days, and their engagement and/or motivational state was not necessarily constant between them. They were sometimes highly engaged (adopting vigorous behavior), while at other times they were less so. To account for such variations, we established a Motivation index that allows comparison across sessions based on the average exploitation duration (ED) and the average speed of the movement (VP). The rationale is that short ED and high VP reflect a higher engagement in the task than longer ED and lower VP, which, in turn, may indicate reduced implicit motivation to decide and act (Mazzoni et al., 2007). To normalize this behavioral variable, we first computed z-scores of average VP and average ED for each monkey. Thus, for each subject, the mean (μ) and standard deviation (σ) of VP and ED across all sessions were used to normalize the values as follows:

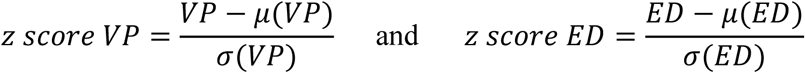

We then calculated the Motivation index for each session of each monkey using the following formula:

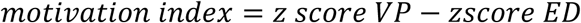

This metric provides a relative measure of animal engagement/motivation at the session level.

### Modeling the optimal decision and movement durations in each trial

One of the objectives of the present study is to determine whether the animals’ behavior, particularly in terms of decision-movement coordination, is close to what would be required to obtain an optimal reward rate on each trial. This optimal reward rate reflects the value of the obtained rewards, discounted by the cost of the upcoming movement effort and the total time needed to obtain them. The reward rate, which therefore depends on both the exploitation (ED) and movement (MD) durations, is denoted as followed:

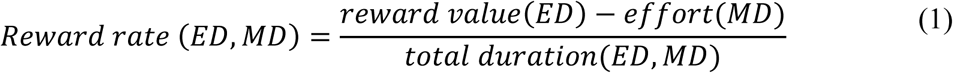

Since each collected point is associated with a drop of juice, their accumulation represents the reward component in equation (1). The accumulation of points over time *t* within each trial (i.e., as a function of the exploitation duration, ED) is given by:

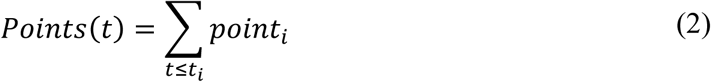

with *i* ranging from 0 to 6 and where *point_i_* corresponds to the point collected at time t*_i_*, which depends on the reward rhythm level (fast, medium or slow).

While the actual delivery of points occurs in a discrete manner, we assume that the underlying process of accumulation over time is gradual and can be approximated by an asymptotic function of the following form:

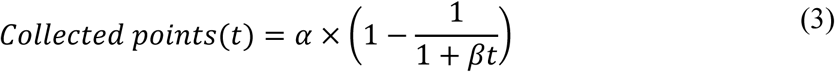

with α = 6 (i.e., the maximum number of points that can be obtained per exploited patch, corresponding to the asymptote of the curve) and β a slope-adjustment parameter fitted to empirical data by minimizing the sum of squared differences between observed values and model predictions. Using MATLAB’s *fminsearch* function, the fitted β values were: 1.37 for the fast rhythm, 0.62 for the medium, and 0.37 for the slow rhythm (Supplementary Figure 1A).

The subjective value of the accumulated rewards is then estimated by using the following temporal discounting function:

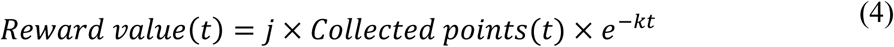

with j = 5 (the subjective value of a single drop of juice, expressed in energy equivalent, i.e., joules) and k = 0.1 (the temporal discounting coefficient). Both parameters were chosen so that the model best predicts the number of rewards accumulated by the animals and the duration of the sessions (Supplementary Figure 2).

The reward value thus depends on both the duration of the exploitation and the point/reward rhythm. It was modeled for a range of ED values between 0 and 6 seconds (Figure 2, top left panel, and Supplementary Figure 1B).

**Figure 2.**
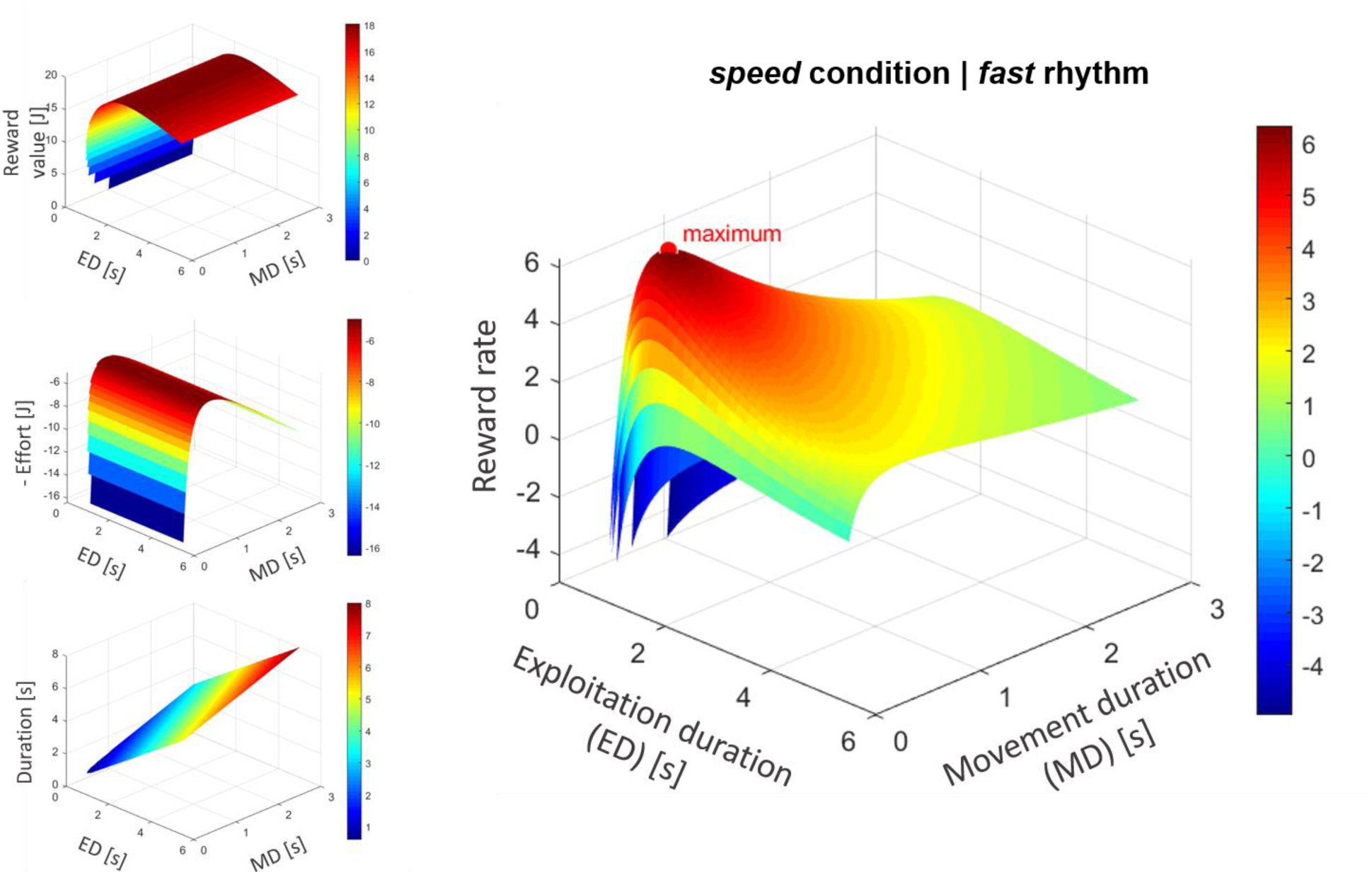
Modeling of reward value, effort, total trial duration, and resulting reward rate for a trial performed in the *fast* reward rhythm and the *Speed* motor condition. The three left panels represent the individual components of the reward rate equation (equation 1 in the main text): reward value (top), movement effort (middle), and total trial duration (bottom). The right panel shows their combination to compute the reward rate as a function of exploitation duration (ED) and movement duration (MD). Reward rate values below -5 are not displayed for visibility. The red dot indicates the maximum reward rate, allowing to extract the optimal exploitation and movement durations (ED_opt and MD_opt).

Next, to model the effort cost associated with performing a reaching movement to leave the current patch, we used the equation proposed by Shadmehr et al. (2016):

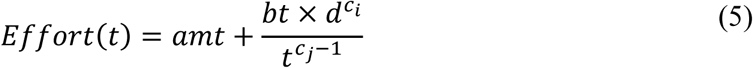

with a = 15, b = 77, cᵢ = 1.1, cⱼ = 2.7; m being the mass of the monkey’s arm, d the movement amplitude (in meters), and t the movement duration (MD, in seconds). The arm mass was estimated based on the monkey’s total body weight (9 kg for monkey G and 6 kg for monkeys B and D) using the following equation: m = body weight × 0.04 - the same proportion as reported for humans (de Leva, 1996).

Motor effort was then computed as a function of MD, across a range from 0.15 to 3 seconds (Figure 2, middle left panel) and for each motor condition. Since movement amplitude was the same between the *Reference* and *Accurate* conditions, and since we did not model the cost of motor control for simplicity, these two conditions produced the same effort function (Supplementary Figure 1C).

Finally, the total trial duration was defined as the sum of ED and MD (Figure 2, bottom left panel).

Reward value, effort and total trial duration were then combined using equation (1) to compute the reward rate for each combination of ED and MD, generating in each case a specific reward rate manifold (Figure 2, right panel and Supplementary Figure 3). The optimal ED and MD values (ED_opt and MD_opt) for each condition were identified as those maximizing the reward rate on this manifold and are thus considered as the local optimal behavioral values for a given trial. As a result, each of the nine combinations of rhythm level and motor condition generates a unique reward rate manifold and thus a distinct optimal reward rate (RR_opt) with corresponding ED_opt and MD_opt values.

Since *Reference* and *Accurate* conditions do not differ in movement amplitude (and effort only depends on movement amplitude in our model), they yield identical reward rate functions. Consequently, the model produces six unique optimal triplets (RR_opt, ED_opt, MD_opt) across combinations (Figure 6A).

### Simulation of an optimal dataset

From a theoretical perspective, we assume that a session is optimal if, for each trial, the behavior adopted maximizes the reward rate, as previously defined. To assess this, we simulated, for each “real” session performed by the monkeys, an “ideal” session consisting of the same number of trials and the same trial types, but in which each trial was performed using the optimal strategy. Because each trial is characterized by a specific combination of rhythm level and motor condition, we could assign the corresponding optimal exploitation duration (ED_opt) and optimal movement duration (MD_opt) derived from the previously modeled manifold.

Then, using the value of MD_opt and the amplitude of the movement (d), we estimated the optimal peak of velocity of the movement (VP_opt) using the following formula:

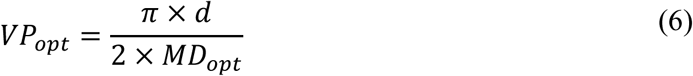

Additional metrics were computed to assess whether monkeys’ performance approached or diverged from theoretical optimality over time. First, we estimated the number of points that would have been collected if the monkeys had always decided to leave the patches at ED_opt, given the rhythm level of each trial. We then calculated the duration of each simulated trial by summing ED_opt and MD_opt plus the fixed holding time of 100ms necessary to start the exploitation phase. Finally, we computed an optimal intake rate (IR_opt) for each simulated session, defined as the total number of collected points divided by the sum of all simulated trial durations. These optimal intake rates were compared to the real intake rate of the corresponding sessions, and we examined how the difference between these two indicators evolved across sessions for each monkey.

### Statistics

We used MATLAB (MathWorks^®^) for all statistical analyses and the significance level was set at 0.05. One-way ANOVAs followed by post-hoc comparisons using Tukey’s test were conducted to examine overall behavioral differences in exploitation duration (ED), movement velocity peak (VP) and intake rates (IR) between the three subjects. Pearson correlations were computed to assess the relationship between ED and VP at the single-trial level (i.e. the degree of coordination between ED and VP, referred as *Rcoeff* hereafter) and at the session-by-session level, both in real and simulated data. Wilcoxon signed-rank tests were used to assess the difference in behavior between real sessions (ED and VP) and simulated ones (ED_opt and VP_opt). Pearson correlations were also used to test the evolution of the intake rate (IR) over sessions, the evolution of the distance between IR and IR_opt across sessions and the session-by-session relationship between Motivation index and IR.

To identify which parameters at the session-level were associated with variations in the intake rate of the optimal simulated data and their relative weight, we performed linear mixed-effects modeling. *Intake rate* (defined as the average number of points collected per unit time within a session) was set as the dependent variable. Four fixed effects were included: *Rcoeff* (the regression coefficient representing the coordination strategy employed), *Motivation index* (the measure of session-level engagement/motivation), *Session* (the order of the sessions) and *Number of trials* (number of trials performed per session, proxy of the number of executed movements). We expected a significant relationship between *Rcoeff* and *Intake rate*, as the coordination between decision and movement is hypothesized to contribute to performance (see details in the Results section). The *Motivation index* was also expected to correlate with *Intake rate*, since higher engagement typically enhances efficiency. In addition, we predicted an effect of *Session* due to the animals’ increasing experience with the task even if this effect is expected to be small since the same strategy was applied across simulated sessions. Finally, the *Number of trials* was also expected to influence performance, as it varies across simulated sessions (being constrained by the corresponding real sessions and thus dependent on each monkey’s behavior). To account for potential inter-individual variability, *Subject* was included as a random effect. Model estimation was carried out using restricted maximum likelihood (REML) in MATLAB (*fitglme* function). Competing models were evaluated based on information criteria (AIC/BIC) and convergence diagnostics, and the final selected model was: ***Intake rate ∼ Rcoeff + Motivation index + Session + Number of trials + (1 | Subject)***. See Supplementary Table 1 for details on the tested models and model comparisons.

## Results

### General observations

On average, monkey G collected 1092 ± 16 drops of juice/points (mean ± SEM) during the 84 sessions she completed, which required 323 ± 5 trials per session, with sessions lasting 28 ± 1 minutes. Monkey B collected 1570 ± 59 points per session during 50 sessions, which required completing 562 ± 21 trials over 31 ± 1 minutes, while monkey D collected 1332 ± 69 points through 552 ± 28 trials in 36 sessions, with sessions lasting 39 ± 2 minutes (Supplementary Figure 2A-C).

Then, we examined overall behavioral differences in exploitation duration (ED), movement velocity peak (VP) and intake rates (IR) across the three subjects (Figure 3). Monkey G exhibited the longest ED (3.49 ± 0.07 s) compared to monkeys B and D (1.75 ± 0.03s and 1.65 ± 0.04s, respectively) (one-way ANOVA, Subject, F_(2, 167)_ = 275, p < 0.001). Post-hoc comparisons using Tukey’s test show that monkey G spent significantly (p < 0.001) more time in the patches than both monkey B and monkey D and that monkey B also exploited the patches significantly longer than monkey D (p < 0.001).

**Figure 3.**
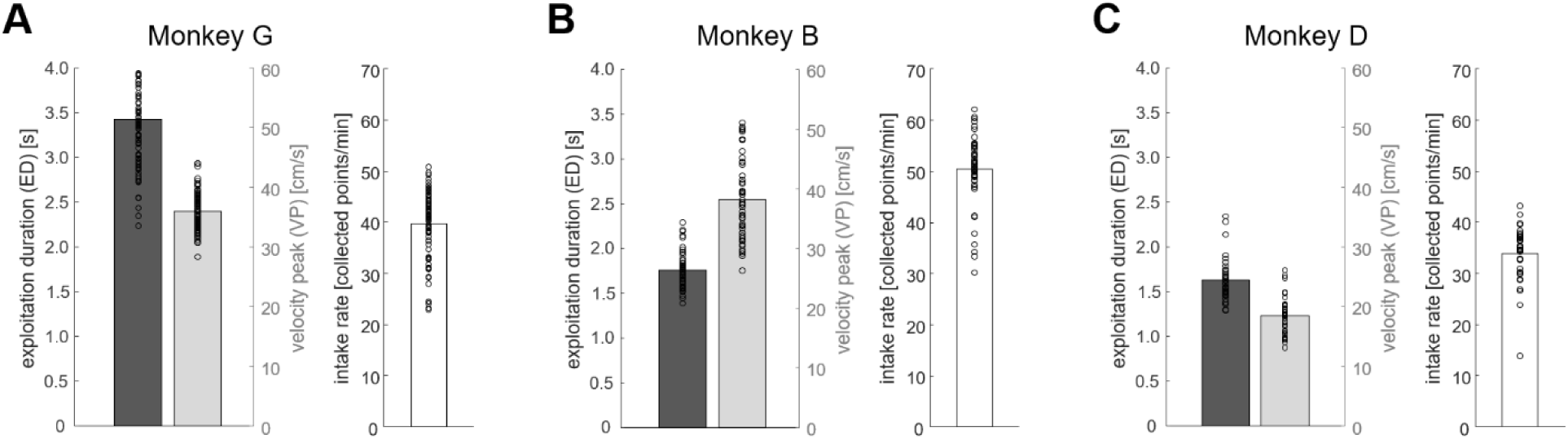
General behavior across sessions. **A**: Exploitation duration (dark gray), reaching movement speed (light gray) and intake rate (number of collected points per minute, white) for monkey G. Bars illustrate mean values across sessions and dots illustrate individual values for each session. **B**: Same as A for sessions performed by monkey B. **C**: Same as A for sessions performed by monkey D.

Monkey B showed the highest VP (38.2 ± 1.0 cm/s), followed by monkey G (36.0 ± 0.4 cm/s) and monkey D (18.5 ± 0.5 cm/s) (one-way ANOVA, Subject, F_(2, 167)_ = 228, p < 0.001). Post-hoc analyses indicate that monkey G executed slower movements than monkey B (p < 0.05) but faster than monkey D (p < 0.001). Monkey B also exhibited significantly higher VPs than monkey D (p < 0.001).

Monkey B exhibited the highest IR (50.3 ± 1.0 points/min), followed by monkey G (39.7 ± 0.7 points/min) and monkey D (33.9 ± 0.9 points/min) (one-way ANOVA, Subject, F_(2, 167)_ = 75, p < 0.001). Post-hoc comparisons confirm that monkey B was overall more efficient in the task than both monkey G (p < 0.001) and monkey D (p < 0.001) and that monkey G’s IR was also significantly higher than that of monkey D (p < 0.001).

### Coordination between decision and movement

Here we examine whether and how the three monkeys coordinated the duration of their exploitations (ED) with the vigor of the arm movements following these exploitations (velocity peak of the reaching movement, VP).

For monkey G, trial-by-trial analyses indicate a significant and positive correlation between ED and VP for the majority of sessions performed (Pearson correlations, p < 0.05 in 48 of 84 sessions). This positive relationship means that longer exploitation durations were followed by more vigorous movements (the “compensation” strategy mentioned in the Introduction). However, in 6 of the 84 sessions, a significant negative correlation is observed, indicative of a co-regulation of decisions and movements (i.e., the longest the exploitations, the slowest the movements, and vice versa). Interestingly, this correlation coefficient overall decreases over the time course of monkey’s experience (Figure 4A, left panel), indicating a compensation strategy that vanishes over time (Pearson correlation across sessions: r = -0.36, p < 0.001).

**Figure 4.**
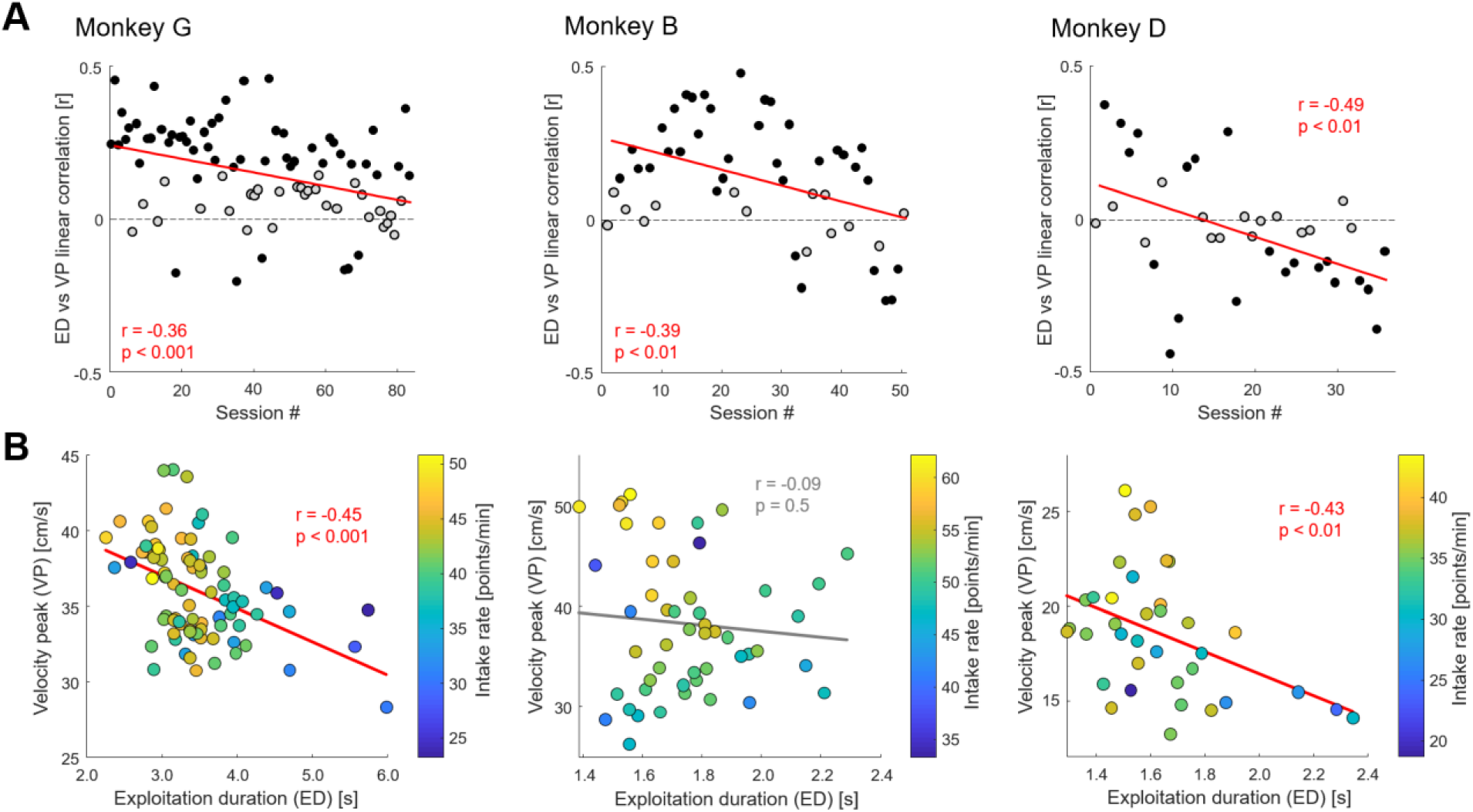
Relation between exploitation duration and arm movement speed within and between sessions. **A**: Trial-by-trial Pearson linear correlation between exploitation duration (ED) and arm movement peak velocity (VP) within sessions performed by monkey G (left panel), monkey B (middle panel) and monkey D (right panel). Each dot illustrates the result of a single session. A negative (or positive) r value indicates that longer exploitations are associated with slower (or faster) movements. Black-filled dots indicate sessions where the Pearson correlation is statistically significant and the red line shows the result of a significant linear regression across sessions. **B**: Relationship between means of exploitation duration (ED) and arm movement peak velocity (VP) across sessions for monkey G (left panel), monkey B (middle panel) and monkey D (right panel). Each dot represents a single session and colors represent within-subject intake rate levels (warmer colors for higher IR). A gray line shows the result of a non-significant Pearson linear regression through the data while a red line shows the result of a significant one.

For monkey B, a significant and positive correlation between ED and VP is observed in 30 out of 50 sessions, also consistent with a compensation mode of coordination between decisions and movements. As for monkey G, this relationship overall decreases over time, as evidenced by a significant reduction in r values across sessions (r = -0.39, p < 0.01, Figure 4A, middle panel). Notably, 6 sessions also exhibit a significant negative correlation, indicative of a co-regulation of decisions and movements in the last sessions performed by monkey B.

Lastly, we observed the same trend for monkey D. The correlation between ED and VP strongly decreases over sessions (r = -0.49, p < 0.01, Figure 4A, right panel), indicating a progressive shift (from a compensation to a co-regulation mode) in the strategy coordinating decision and action processes. A significant positive correlation is observed in 7 out of 36 sessions, while a significant negative correlation is shown in 14 sessions, mostly during the last sessions performed by monkey D.

Next, we investigated the relationship between ED and VP not at the individual trial level as we have done so far but more globally, at the session level.

For monkey G, a significant negative correlation is observed between sessions (Figure 4B, left panel; r = -0.45, p < 0.001), indicating that in sessions during which she made the shortest exploitations, she also executed the more vigorous movements. A similar pattern was found in monkey D (Figure 4B, right panel; r = -0.43, p < 0.01). By contrast, no significant correlation between ED and VP was found at the session level for monkey B (Figure 4B, middle panel; r = - 0.09, p = 0.5). Another important result is that sessions with a higher Motivation index are also those with higher IR (r = 0.51, p < 0.001 for monkey G; r = 0.42, p < 0.01 for monkey B; and r = 0.52, p < 0.01 for monkey D). This relationship also appears in Figure 4B, where warmer-colored dots (i.e., sessions with higher IR) broadly correspond to sessions characterized by shorter ED and higher VP.

### Comparison between observed monkey performance and simulated optimal performance

In a simulated dataset, we generated for each session performed by the monkeys an optimal session consisting of the same number of trials, but in which each trial was performed maximizing the local reward rate (see the Supplementary information section of the article for details on the simulated mean ED and mean VP for each monkey).

We examined the evolution of the intake rate (IR) across the monkeys’ different sessions, defined for each session as the number of points collected per minute of task performed. It is important to note that this variable does not account for the impact of motor effort. A linear regression of these data shows that IR increases across sessions for both monkey G and monkey B (Figures 5A and 5B left panels; Pearson correlations: r = 0.41, p < 0.001 for monkey G; r = 0.38, p < 0.01 for monkey B). A similar trend is observed for monkey D (Figure 5C left panel; r = 0.29, p = 0.09).

**Figure 5.**
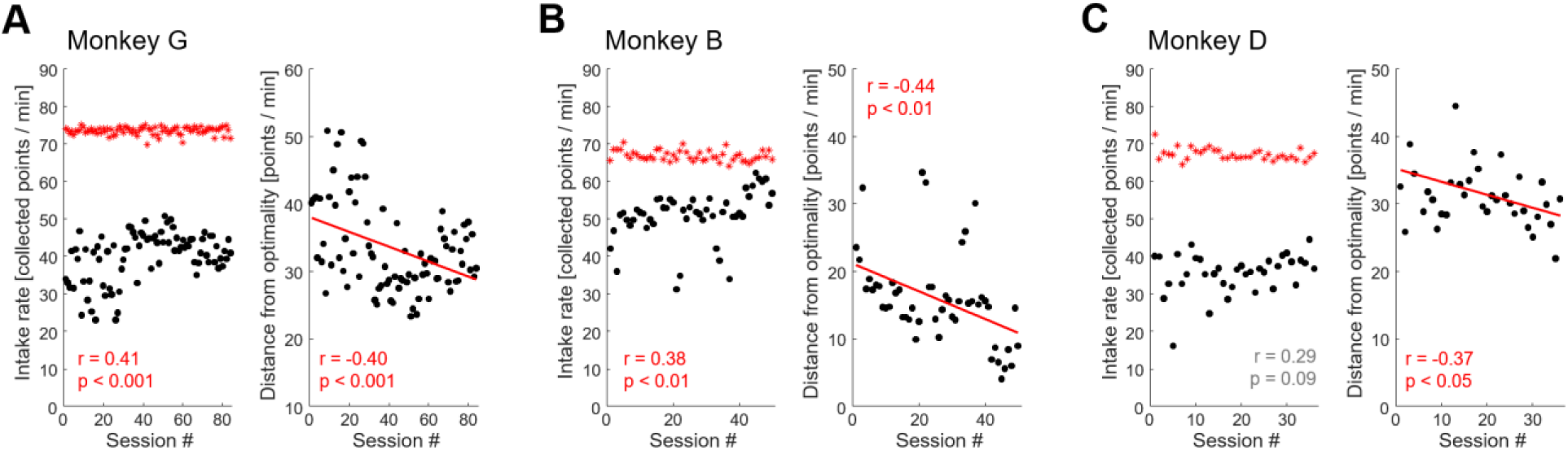
Evolution of intake rates across sessions and distance from optimality. **A**: *Left panel*. Within-session intake rate, defined as the number of points collected per minute, as a function of sessions performed by monkey G. Black-filled dots illustrate “real” intake rates and red asterisks show the simulated optimal intake rates for each corresponding simulated session. *Right Panel*. Evolution of the distance, in points per minute, between real and simulated intake rates across sessions. The red line shows the result of a significant linear regression across sessions. **B**: Same as A for sessions performed by monkey B. **C**: Same as A for sessions performed by monkey D.

**Figure 6.**
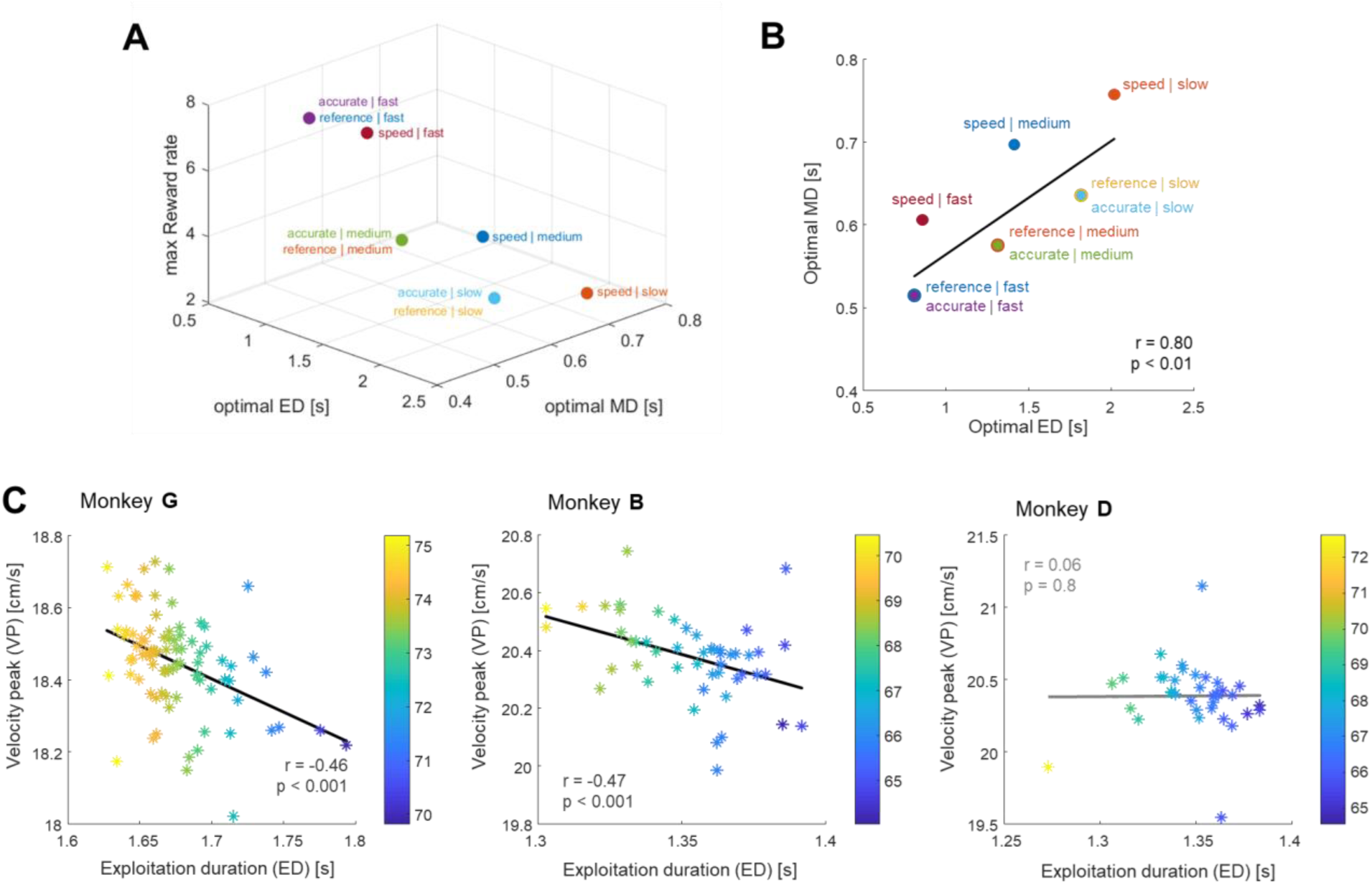
Optimal behavior across combinations of reward rhythm and motor condition and the associated coordination strategy within and between simulated sessions. **A**: For each of the 9 combinations of rhythm levels and motor conditions, we display the triplet of values corresponding to the optimal reward rate, along with the associated optimal exploitation duration (ED_opt) and movement duration (MD_opt). Data are based on modeling with j = 5, k = 0.1 and m = 6kg in equations (4) and (5). **B**: Orthogonal projection of these 9 combinations in the ED-MD space. The black line represents the result of a significant positive linear regression through the data, indicating that a co-regulation of decisions and actions allows to maximize the rate of reward. **C**: Pearson linear correlation between means of optimal exploitation duration (ED_opt) and arm movement peak velocity (VP_opt) in simulated data for monkey G (left panel), monkey B (middle panel) and monkey D (right panel). Each asterisk represents one simulated session and colors represent within-subject simulated intake rate levels (in points/min). A gray line shows the result of a non-significant linear regression through the data while a black line, a significant one.

When considering the simulated optimal sessions, the optimal IR remains stable across time for monkey G (r = 0.01, p = 0.9) whereas it showed a slight decline for monkeys B (r = -0.27, p = 0.05) and D (r = -0.37, p < 0.05). These small variations probably reflect differences in the proportion of trial types experienced and in the overall number of trials performed by each animal. Most notably, the difference, or distance, between real and optimal IR decreases over sessions for all animals, indicating a progressive convergence toward this optimal behavior, mainly driven by the increase in real IR (Figure 5A-C, right panels; Pearson correlations r = -0.40, p < 0.001 for monkey G; r = -0.44, p < 0.01 for monkey B; r = -0.37, p < 0.05 for monkey D).

### Coordination between decision and action in the optimal simulated dataset

In the following paragraphs, we investigate whether the optimal strategy, i.e. the behavior maximizing the reward rate on a trial-by-trial basis, relies, at least partly, on a specific coordination between decision and movement processes.

To do so, we first projected the optimal triplets (reward rate, ED, MD) for each of the nine combinations of rhythm level and cost condition (Figure 6A and Supplementary Figure 3) onto the two-dimensional ED_opt-MD_opt space (Figure 6B). This orthogonal projection shows a significant positive correlation between the optimal exploitation (ED_opt) and movement (MD_opt) durations across the different task conditions (r = 0.80, p < 0.01 for a monkey weighing 6kg; r = 0.76, p < 0.05 for a 9kg one). This observation suggests that longer exploitations are associated with longer movements in this simulation of optimal behavior, indicating that a co-regulation of decisions and movements allows reward rate maximization.

As performed on the real dataset, we then assessed the decision-movement coordination mode (i.e. no correlation, co-regulation or compensation) in the simulated sessions, by computing the trial-by-trial Pearson correlation coefficient between the optimal exploitation duration (ED_opt) and the optimal velocity peak (VP_opt) for each simulated session. We found in all simulated sessions and for the three animals a significant negative correlation between ED_opt and VP_opt (r = -0.56 ± 0.004 across sessions for monkey G; r = -0.64 ± 0.003 for monkey B; r = -0.65 ± 0.006 for monkey D), indicating a strong and robust co-regulation mode of coordination in which shorter exploitations are associated with faster movements, and vice versa. Additionally, no evolution of these correlation coefficients across sessions is observed, supporting a stable coordination pattern in this optimal behavior.

When examining the coordination between the optimal exploitation duration and peak velocity at the session level in the simulated data, results similar to those observed in the real dataset emerge. No apparent relationship was found between average ED_opt and average VP_opt for monkey D (Figure 6C, right panel; r = 0.06, p = 0.8) but, for monkeys G and B, a significant negative correlation was found (Figure 6C, left and middle panels; r = -0.46, p < 0.001 for monkey G; r = - 0.47, p < 0.001 for monkey B), indicative of a co-regulation of exploitations and movements at the session level in the simulated dataset.

Finally, to assess which factors best explained the intake rate (*IR*, expressed in points/min) in optimal simulated data, we fitted a linear mixed-effects model including *Rcoeff* (coordination strategy), *Motivation index* (session-level engagement/motivation), *Session* number and *Number of trials* as fixed effects and *Subject* as a random intercept. The model provided a good fit to the data and revealed significant effects of all four predictors on *IR* (Table 2). The effects of *Session* (β = -0.013, p < 0.001) and *Number of trials* (β = -0.001, p < 0.05) were relatively small. By contrast, *Rcoeff* had a large (the largest of all factors tested) negative effect on *IR* (β = -11.87, p < 0.001), indicating that if *Rcoeff* decreases by one unit (i.e., more co-regulation), this is associated with an increase of 11.87 points/min in *IR*. The *Motivation index* was a positive predictor (β = 0.60, p < 0.001), with higher values corresponding to higher *IR*.

**Table 2.** Fixed effects estimates from the selected linear mixed-effects model. For each parameter, the estimated coefficient (β), standard error (SE), t-statistic (tStat), degrees of freedom (DF) and p-value are reported.

| Name | Estimate ($\beta$) | SE | tStat | DF | p value |
| --- | --- | --- | --- | --- | --- |
| Intercept | 63.26 | 2.31 | 27.44 | 165 | < .001 |
| *Session* | -0.013 | 0.0027 | -4.73 | 165 | < .001 |
| *Number of trials* | -0.001 | 0.0005 | -2.03 | 165 | < .05 |
| *Rcoeff* | -11.87 | 1.489 | -7.97 | 165 | < .001 |
| *Motivation index* | 0.60 | 0.031 | 19.29 | 165 | < .001 |

## Discussion

In this study, we investigate whether and how monkeys coordinate decisions and subsequent movements during foraging and, if so, whether this coordination supports trial-by-trial reward rate maximization. Results show that in most sessions, the three animals coordinated the duration of their exploitations with the vigor of their movements and that their coordination strategy evolved with experience in the task from a compensation to a co-regulation mode. This evolution coincides with an increase of their intake rates, reflecting higher foraging performance over sessions. Then, we simulated optimal behavior that maximized reward rate on a trial-by-trial basis. Although these optimal data indicate intake rates higher than those achieved by monkeys, the difference between observed and simulated rates systematically and consistently decreases with the animals’ experience, strengthening the possibility that monkeys gradually converged toward this optimal behavior. Finally, we assessed from the theoretical data the type and the weight of the decision-movement coordination in foraging performance. The co-regulation of decisions and movements univocally emerged for all monkeys and the linear mixed-effects model showed that this coordination has a very large influence on foraging efficiency.

These findings extend previous evidence on decision-movement coordination strategies, and their evolution with experience in the task. In a perceptual decision-making task, we recently reported that monkeys shifted from a co-regulation to a compensation strategy over sessions (Saleri and Thura, 2024). In that context, humans and monkeys usually compensate for the temporal cost of long deliberations through more vigorous movements (Thura et al., 2014; Thura, 2020; Saleri Lunazzi et al., 2021, 2023; Leroy et al., 2025) in order to be more efficient (i.e. to optimize the number of correct trials per minute). A similar tendency was observed in the present data (see Supplementary Figure 4) as sessions with longer exploitation durations were also those in which monkeys relied more on compensation strategy (or co-regulated less). Nevertheless, the shift in strategy across sessions was reversed in the present foraging task compared to the studies mentioned above in which subjects performed a perceptual decision-making task. One possible explanation is that the three monkeys have been initially trained in the perceptual decision-making task for which compensation was more advantageous (among the three animals, two were the subjects of the study described in Saleri and Thura, 2024). As a result, they may have initially applied the same strategy before progressively shifting toward co-regulation. Indeed, unlike many perceptual decision tasks, the present foraging task does not impose strong temporal constraints which reduces the need for compensation. In addition, accuracy is not an important factor in this task since no errors can occur. Finally, since foraging tasks are generally very intuitive and therefore simple to learn and perform, calling upon the natural behavioral repertoire of all animals (Calhoun and Hayden, 2015), it is possible that co-regulation, proposed to involve simple and energy efficient neural mechanisms (Carland et al., 2019; Thura et al., 2025), is naturally favored and selected by subjects as a default way to coordinate decisions and movements (Fievez et al., 2024). Together, these differences between tasks highlight the flexibility of decision-movement coordination mechanisms, which adapt to task paradigms.

We also found that, at the session level, the coordination between exploitation duration and movement vigor in observed (and simulated) data point to a co-regulation strategy. This observation may reflect a global, non-specific modulation of the animal’s motivational state across sessions, consistently with the recently reported observations suggesting that decision-modulating signals, notably urgency, have a wide impact on neural responses, modulating multiple brain regions both at the cortical and subcortical levels (Murphy et al., 2016; Steinemann et al., 2018; Carland et al., 2019; Kaduk et al., 2023). To evaluate this more formally, we computed a Motivation index that includes both mean movement vigor and mean exploitation duration for each session and found that it also positively co-varies with the performance of animals during foraging. Interestingly, a high Motivation index, defined as short exploitation duration and high movement vigor, may also reflect impulsivity, particularly for monkeys B and D who are younger than monkey G (Green et al., 1999). Further work is needed to better define the contours of optimal behavior, namely where impulsivity begins and where optimality consequently ends, highlighting the complexity of decision-action coordination mechanisms which operate at multiple scales.

Despite their improvement in efficiency across sessions, our monkeys remained sub-optimal relative to the simulated optimal behavior. Several factors may explain this. First, animals frequently disengaged from the task, leading to longer session durations compared to the simulations where trials run consecutively. Second, empirical exploitation durations were consistently longer than optimal, indicative of overharvesting strategies, as reported in other foraging contexts (Cash-Padgett and Hayden, 2020). Third, movements of monkeys G and B were more vigorous than those simulated, potentially increasing energetic costs and thereby devaluing reward value (Shadmehr et al., 2016). By contrast, monkey D exhibited slower movements, minimizing energetic expenditure but resulting in longer trial durations (Myerson and Green, 1995).

Moreover, although we believe that the present work convincingly indicates that optimal behavior during foraging requires the co-regulation of exploitations and movements, several limitations and perspectives to this initial work must be mentioned. First, the simulations did not take into account the requirement imposed on monkeys to collect a fixed number of points to complete a session. Moreover, even if the number of simulated trials was matched to the real sessions, the model did not reach the fixed reward goal required from the monkeys. As a result, the simulated sessions yielded fewer collected points than the task goal (Supplementary Figure 3). If the model had been constrained to reach it, more trials would have been necessary, increasing energy expenditure. One perspective would also be to incorporate variability into the simulated data in order to better approximate real behavior. Finally, and most importantly, our theoretical work on behavioral optimality is based on the assumption that animals seek to maximize their reward rate at a very local scale, during each individual trial. However, it is more than likely that other factors come into play and modulate how optimal behavior should be estimated. For example, current theories of foraging such as the Marginal Value Theorem states that to maximize the global utility, the animal should compare the local intake rate in the current patch with the average rate available in the environment, namely the global intake rate (Charnov, 1976; Yoon et al., 2018). Still, we interpret monkeys’ behavior in the present study as a progressive adaptation of their coordination strategy as they learn the task structure and environmental contingencies and adapt their behavior in consequence. For the simulated data, a promising perspective will be to update the model in order to assess whether the predicted optimal behavior under MVT assumptions leads to comparable coordination strategies.

In conclusion, we provide novel evidence that monkeys flexibly coordinate decision and action in a foraging paradigm, and that such coordination progressively approaches, but does not achieve, simulated optimal performance at a local scale. This work contributes to demonstrate that coordination strategies are highly context-dependent and it opens perspectives for investigating how animals integrate environmental contingencies to adjust coordination strategies at more global scales.

## Authors’ contribution

CS and DT designed the experiment and coded the task

GG and CS trained the monkeys and collected the data

CS and AF conducted the analyses and prepared the figures

CS and DT wrote the draft of the manuscript

CS, GG, AF and DT revised the draft and approved the final version of the manuscript

## Data availability statement

The datasets used and/or analyzed during the current study are available from the corresponding authors on reasonable request.

## Acknowledgements/Funding

The authors thank Sonia Alouche for her administrative assistance, Frédéric Volland for his significant contribution during the technical preparation and maintenance of this experiment, Michel Filiptchenko and Manon Dirheimer for their daily care of the monkeys in our animal facility. They also thank Nils Kolling for help on data modeling and Luc Renaud who performed most of the surgical procedures mentioned in this study.

This work is supported by an ANR grant (ANR-22-CE37-0010-02/ BasalCost) to DT.

