## Supplemental information for "Optimal foraging requires coordinating decisions with movements"

*Supplementary Information*

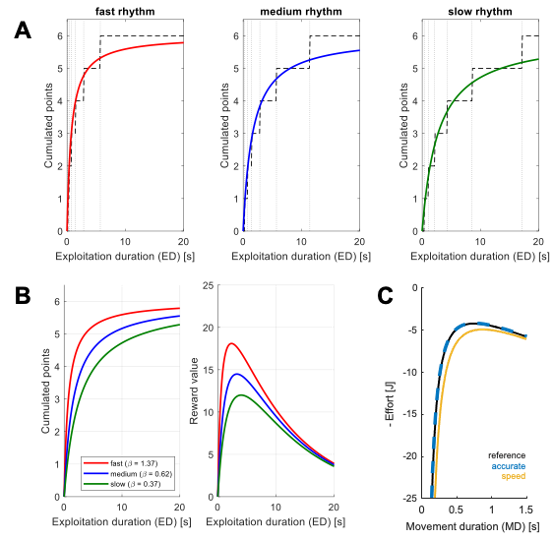

**Figure S1 - Modeling of Reward and Effort functions.** **A**: Empirical and modeled reward rhythms. Vertical dotted lines mark the timing of the six potential juice drops obtainable by holding the cursor in a patch. Dashed lines represent the actual reward intake function, while colored curves show the modeled proxies (equation 3 in the main text; slow in green, medium in blue and fast in red). **B**, Left panel: Models of the three reward intakes function. See details in Materials and Methods. β parameters were estimated with MATLAB’s *fminsearch* (1.37 for the fast rhythm, 0.62 for the medium, and 0.37 for the slow rhythm). Right panel: Subjective value of accumulated rewards estimated with a temporal discounting function (equation 4 in the main text, data are based on modeling with j = 5 and k = 0.1). **C**: Effort cost of reaching movement in the three different motor conditions, computed with the equation from Shadmehr et al. (2016) (equation 5 in the main text). Data are based on modeling with m = 6 kg and the reference condition is represented in black, the accurate condition in blue and the speed condition in yellow.

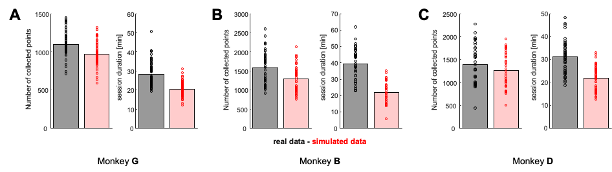
**Figure S2 - Number of collected points per session and session durations in real and simulated data.** **A**, Left panel: Number of collected points in real sessions (dark gray) and in optimal simulated sessions (red). Bars illustrate mean values across sessions and dots illustrate individual values for each session. Right panel: Corresponding session durations. **B**: Same as in A for sessions of monkey B. **C**: Same as in A for sessions of monkey D.

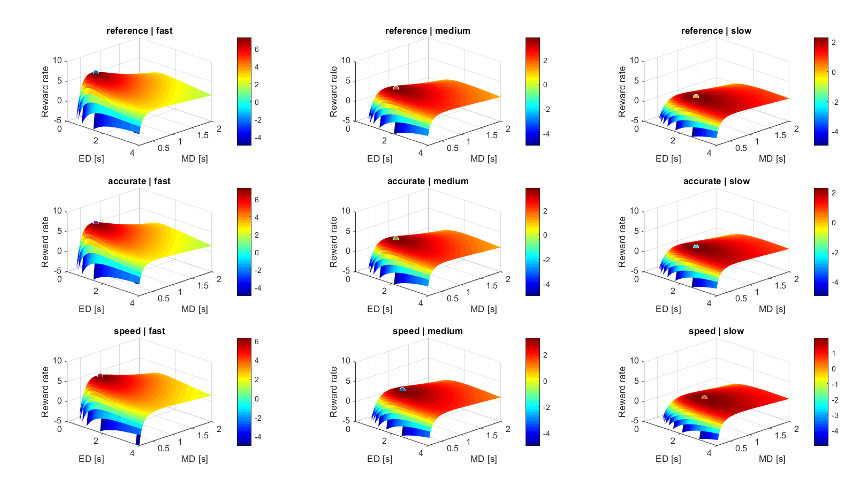
**Figure S3 - Optimal behavior across the nine combinations of reward rhythm and cost conditions.** Manifolds of reward rates of the 9 combinations of rhythm levels and cost conditions (see equation 1 in the main text). Reward rate values below -5 are not displayed for visibility. The colored dots indicate the maximum reward rate of each combination, allowing to extract the optimal exploitation and movement durations (ED_opt and MD_opt). Data are based on modeling with j = 5, k = 0.1 and m = 6 kg.

Comparison between observed monkey performance and simulated optimal performance

As mentioned in the main text, we generated for each session performed by the monkeys an optimal session consisting of the same number of trials, but in which each trial was performed maximizing the local reward rate. Results indicate that exploitation durations (ED) are shorter in the simulated data than in the real data (mean ± SEM = 1.67 ± 0.003 s, p < 0.001 for monkey G, 1.35 ± 0.003 s, p < 0.001 for monkey B and 1.35 ± 0.004 s, p < 0.001 for monkey D). By contrast, simulated movement vigor (VP) is lower than the real data for monkeys G and B (18.4 ± 0.01 cm/s, p < 0.001 for monkey G; 20.4 ± 0.02 cm/s, p < 0.001 for monkey B), whereas monkey D was slower compared to the simulated data (20.4 ± 0.04 cm/s, p < 0.01).

In order to investigate whether the mode of coordination (i.e., the sign of the correlation between ED and VP) was related to the average ED across sessions (as shown in Saleri and Thura, 2024), we computed Pearson correlations between this R coefficient and the average exploitation duration across sessions, for each subject and for both real and simulated data.

For monkey G, the coordination mode does not appear to be related to the session’s average ED (Supplementary Figure 4, top left panel; r = -0.09, p = 0.4). By contrast, for monkeys B and D, a significant positive correlation is observed between ED and the ED-VP correlation between sessions (Supplementary Figures 4, middle and right panels; r = 0.47, p < 0.01 for monkey B and r = 0.42, p < 0.01 for monkey D). This observation means that sessions with longer ED are associated with stronger compensatory strategies of decision and movement durations (i.e. higher values of r). In other words, the longer the ED, the more the monkeys compensate for this long exploitation with a short movement.

Then, we examined this same relationship in the simulated data. For the simulated datasets of monkeys G and D, a significant positive correlation was observed between mean ED_opt and the ED_opt-VP_opt correlation coefficients (Supplementary Figure 4, bottom left and right panels; r = 0.41, p < 0.001 for monkey G; r = 0.36, p < 0.05 for monkey D). For the simulated trials faced by monkey B, the analysis reveals the same trend without reaching the significance level (Supplementary Figure 4, bottom middle panel; r = 0.26, p = 0.07) This indicates that longer exploitation durations were associated with weaker co-regulation of exploitation duration and movement velocity (i.e., less negative r values).

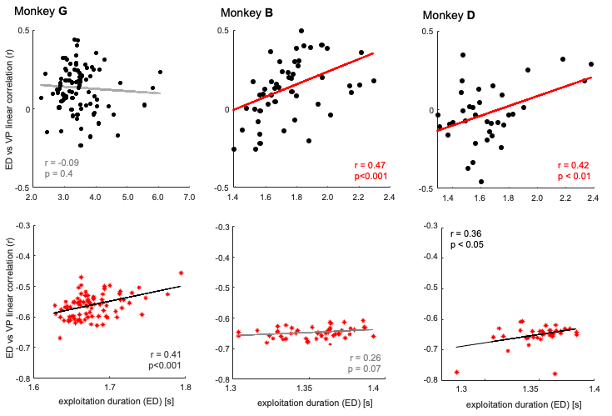
**Figure S4 - Relation between average exploitation duration and coordination mode in real and simulated data**. Top panels: Relationship between Pearson r values (ED-VP linear correlation) and the mean exploitation durations for real sessions. Each dot represents a single session. A gray line shows the result of a non-significant linear regression through the data while a red line shows the result of a significant one. Bottom panels: Same analyses for simulated sessions. Each red asterisk represents one optimal simulated session. A gray line shows the result of a non-significant linear regression through the data while a black line, a significant one.

| **Tested models** | **AIC** | **BIC** | **Log-likelihood** | **p value** |
| --- | --- | --- | --- | --- |
| IntakeRate ~ Rcoeff | 818.8 | 828.2 | -406.4 |  |
| IntakeRate ~ Rcoeff + MotivationIndex | 792.0 | 804.6 | -392.1 | < .001 |
| IntakeRate ~ Rcoeff + MotivationIndex + Session | 753.0 | 768.7 | -371.5 | < .001 |
| IntakeRate ~ Rcoeff + MotivationIndex + Session + nb_Trials | 669.4 | 688.2 | -328.7 | < .001 |
| IntakeRate ~ Rcoeff + MotivationIndex + Session + nb_Trials + (1 \| Subject) | 371.0 | 393.0 | -178.5 | < .001 |

**Table S1 - Tested linear mixed-effects models and model comparison for simulated data.** Each row corresponds to a tested model with different fixed effects. For each model, Akaike Information Criterion (AIC), Bayesian Information Criterion (BIC) and log-likelihood values are reported. The last column indicates the p-value obtained from likelihood ratio tests (LRT) comparing each model to the preceding one.
